# Artificial intelligence supports automated characterization of differentiated human pluripotent stem cells

**DOI:** 10.1101/2023.01.08.523148

**Authors:** Katarzyna Marzec-Schmidt, Nidal Ghosheh, Sören Richard Stahlschmidt, Barbara Küppers-Munther, Jane Synnergren, Benjamin Ulfenborg

## Abstract

Revolutionary advances in AI and deep learning in recent years have resulted in an upsurge of papers exploring applications within the biomedical field. Within stem cell research, promising results have been reported from analyses of microscopy images to e.g., distinguish between pluripotent stem cells and differentiated cell types derived from stem cells. In this work, we investigated the possibility of using a deep learning model to predict the differentiation stage of pluripotent stem cells undergoing differentiation towards hepatocytes, based on morphological features of cell cultures. We were able to achieve close to perfect classification of images from early and late time points during differentiation, and this aligned very well with the experimental validation of cell identity and function. Our results suggest that deep learning models can distinguish between different cell morphologies, and provide alternative means of semi-automated functional characterization of stem cell cultures.

## Introduction

Developments in artificial intelligence (AI) over the past decade have promoted a surge of applications in different fields of sciences. Importantly, the success of deep learning has revolutionized image analysis, not only in computer vision but also for medical and cellular imaging. In contrast to traditional machine learning techniques, deep learning networks are able to automatically and efficiently learn higher-level representations of data without the need for manual feature engineering (Moen et al., 2019). Within the field of stem cell research, deep learning applied to cellular images holds the potential for accurate and automated analysis of cell cultures, as recent studies have demonstrated (Grafton et al., 2021; Guan et al., 2021; Imamura et al., 2021; Joy et al., 2021; Maddah et al., 2020; Zhang et al., 2021). In the study by Imamura et al., induced pluripotent stem cells (iPSCs) were generated from cells from healthy controls and patients with amyotrophic lateral sclerosis. The iPSCs were differentiated into motor neurons and immunostained, followed by image classification where 90 % of the cell images were correctly classified (Imamura et al., 2021). Maddah et al. investigated structural toxicity of iPSC-derived hepatocytes and cardiomyocytes by treating cells with known toxic and non-toxic compounds. A network model trained on fluorescence microscope images was able to correctly identify structural changes from toxic compounds for both cell types (Maddah et al., 2020). By developing a new ensemble-based deep learning method, Joy et al. were able to identify individual nuclei in dense human iPSC (hiPSC) colonies and perform cell tracking to study cell behavior over time (Joy et al., 2021).

A commonly used deep learning architecture specialized for image classification is the convolutional neural network (CNN). It performs mathematical operations to translate an image of pixels into so-called feature maps, which represent visual features like edges and shapes. Several convolutional layers can be stacked on top of each other, each taking the previous map as input, such that the CNN learns higher-level features that are more informative for image classification (LeCun et al., 2015). Several recent studies have demonstrated the feasibility of using CNNs for predicting stem cell differentiation state based on microscopy images. Waisman et al. differentiated mouse embryonic stem cells (mESCs) into epiblast-like cells, and trained a deep learning model to distinguish between culture images of mESCs and differentiated cells. They achieved perfect classification accuracy after only six hours following onset of differentiation (Waisman et al., 2019). Similarly, Liu et al. differentiated human embryonic stem cells (hESCs) into trophoblast-like cells and trained a model to distinguish between trophoblast morphology and hESCs. Two networks used achieved > 99 % accuracy on day 12 (Liu et al., 2021). In studies by Zhu et al. (Zhu et al., 2021) and Lan et al. (Lan et al., 2021), single-cell images were used instead to successfully recognize neural and osteogenic differentiation, respectively.

The immense expectations on pluripotent stem cells (PSCs) in many areas of biomedical research places high demands on production procedures and rigorous quality control of stem cell cultures, e.g., to identify successfully differentiated cultures, and to verify cell marker expression, cellular morphology and functionality. Since this is routinely done experimentally with microscopic inspections, qPCR, immunocytochemistry and various functional assays, the process is costly, time-consuming and requires highly trained specialists. Hence there is a need for more efficient validation methods of PSC identity and function (Coronnello and Francipane, 2021). If these steps could be automated through deep learning-based image analysis, large reductions in cost and workload required for stem cell production could be made.

A few studies have explored the potential of using deep learning for quality control and automated assessment of stem cell cultures. Orita et al. trained a CNN to distinguish between hiPSC-derived cardiomyocyte cultures based on whether they were judged suitable for downstream experiments (Orita et al., 2019). The model achieved close to 90 % accuracy, demonstrating the possibility of using deep learning for automated hiPSC quality control. In a study by Hirose et al., a deep learning method was developed for automated cell tracking of keratinocyte stem cells (Hirose et al., 2021). The aim was to enable non-invasive and efficient quality control of cell cultures proliferative capacity, as an alternative to single-cell clonal analysis, which is expensive and demanding to perform. By relying on cell motion data from the deep learning method, the authors were able to significantly increase the probability of obtaining proliferative stem cell colonies. Similarly, Piotrowski et al. developed a deep learning method for non-invasive automated cell state recognition of hiPSC colonies (Piotrowski et al., 2021). This method was able to distinguish between different cell states, such as hiPSC colony, differentiated cells and dead cells, more accurately than a human expert.

The aim of the present study is to provide a proof-of-concept that CNNs can be used for automated quality control of cultures undergoing hepatocyte differentiation, and at least partially supplant the need for visual inspection of cultures by an expert, in order to evaluate the maturity of the stem cell-derived hepatocyte cultures. In this study, we hypothesize that the morphological features of cells captured in phase-contrast microscopy images are sufficient to distinguish between early and late stages of hepatocyte differentiation. Late stage corresponds to mature hepatocytes with functional properties that recapitulate many features of primary human hepatocytes (Ghosheh et al., 2020; Holmgren et al., 2020). Once a CNN has been trained to recognize these features, it could provide predictions of differentiation stages in line with data obtained from experimental verification.

## Results

### CNN accurately distinguishes between early and late hepatocyte differentiation stages

To test our hypothesis, 1,331 phase-contrast microscopy images were obtained between days 1 to 23 following onset of differentiation of several human PSC (hPSC) lines towards hPSC-derived hepatocytes (hPSC-HEP). In accordance with the differentiation protocol, the culture medium was changed to a hepatocyte maturation medium on day 14 to direct the differentiation from hepatic progenitors into functional hepatocytes. We wanted to investigate the possibility of distinguishing between images captured before and after this timepoint in the protocol, based on morphological features of the cells. Therefore, images collected during days 1-14 were labeled as Early Differentiation (ED, in total 693 images) and images collected during days 16-23 as Late Differentiation (LD, in total 638 images). No images were collected at day 15. The original images were cut and processed into a larger dataset of 6,972 smaller image patches prior to analysis with CNN.

A CNN architecture was then trained on 5,086 images and validated on 1,272 for hyperparameter tuning. The trained model was subsequently evaluated on the independent test set of 614 images to assess its predictive power. Image classification performance during training, validation and testing is presented in Fig. 1, showing that the converged model was highly accurate. Training set accuracy converged on 0.98 after around 200 epochs and validation set accuracy on 0.95 slightly earlier, albeit with greater variation between epochs. The CNN was likewise accurate on the independent test set, where accuracy and F1 of 0.95 were achieved (Fig. 1C). This corresponded to 592 out of 614 images correctly classified as ED or LD.

**Figure 1.**
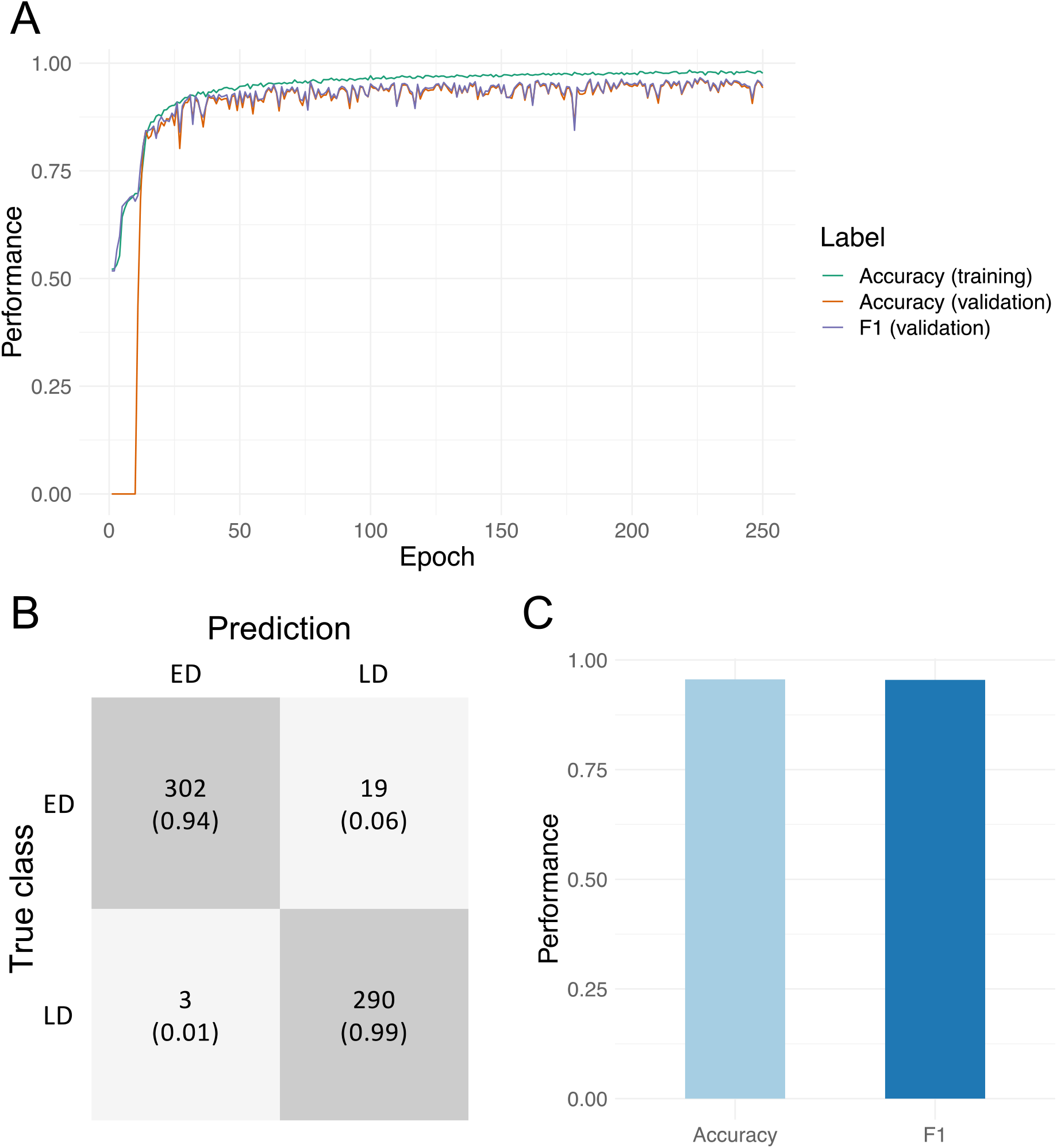
Image classification performance of the CNN. A. Training and validation accuracy and F1. B. Confusion matrix for the independent test set of 614 hiPSC images classified using the trained model. C. Accuracy and F1 obtained for the independent test set. ED: Early differentiation; LD: Late differentiation.

To investigate the relationship between cell culture morphology and predictions made by the CNN, we looked at cell culture images from early and late differentiation, and the reported classification confidence from the CNN. The confidence is computed by the output layer of the model and is a value between 0 and 1, where 0 corresponds to high confidence in ED and 1 to high confidence in LD. This allowed us to compare the morphology in ED and LD images that were correctly and wrongly classified, as well as where the CNN was unsure about the differentiation stage (confidence close to 0.5). Representative images are shown in Fig. 2, with correctly classified in the top left and bottom right (marked by green border), wrongly classified in the top right and bottom left (marked by blue border), and images where the CNN was unsure in the middle.

**Figure 2.**
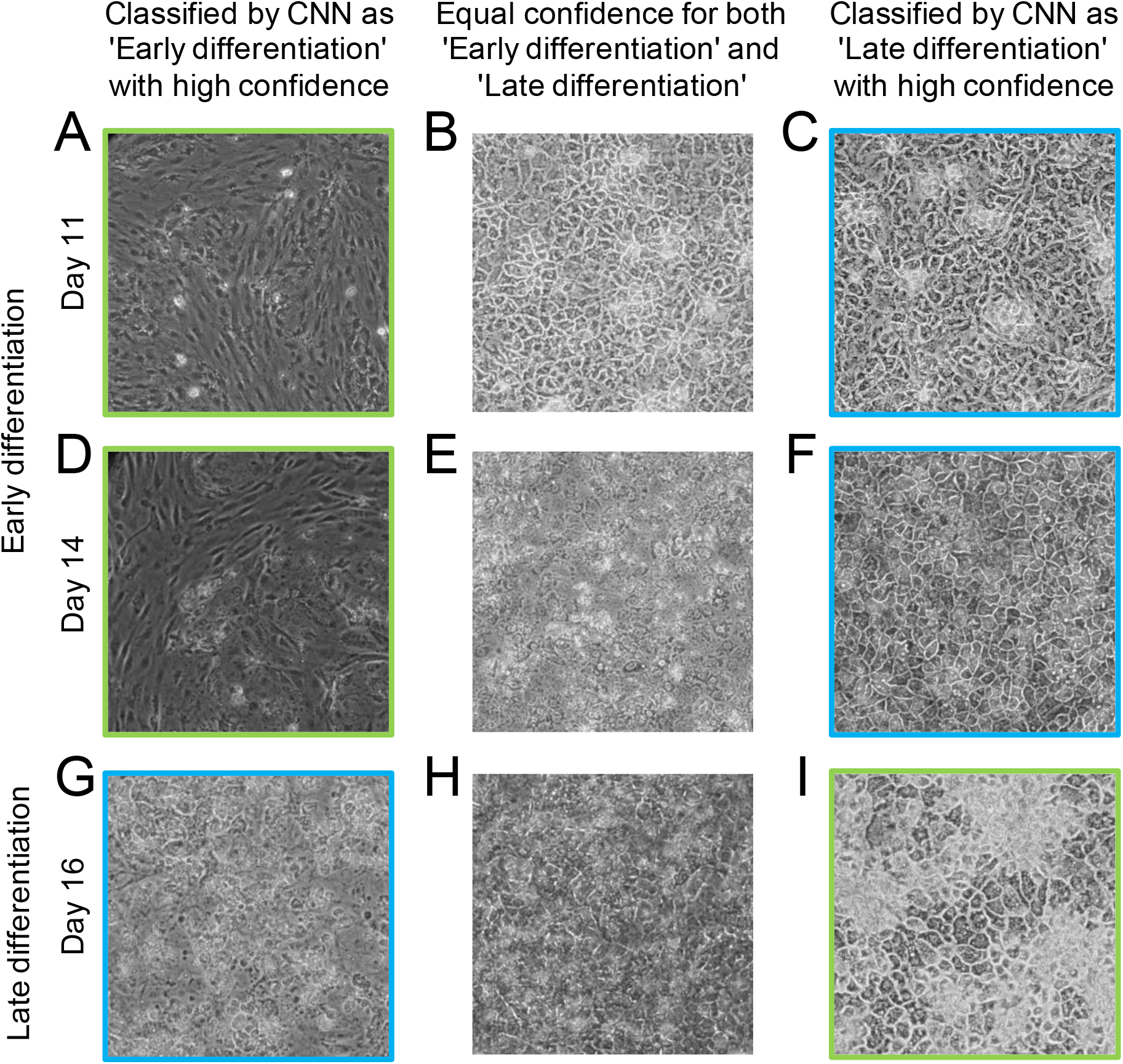
Representative images of hPSC cultures arranged according to their CNN classification. Images correctly classified with high CNN confidence are shown in the left column for Day 11 and Day 14 (panel A and D), and the right column for Day 16 (panel I), marked with green border. hPSC images classified as ED and LD with approximately equal confidence are presented in the middle column (panel B, E and H). Images incorrectly classified with high confidence are shown in the right column for Day 11 and Day 14 (panel C and F) and the left column for Day 16 (panel G), marked with blue border.

A potential explanation for incorrect classification of some images is a similar polygonal cell shape in the progenitor stage and in the hepatocyte stages. Thus, some ED pictures were incorrectly classified by the CNN model as LD, though they do not show the typical hepatocyte morphology (Fig. 2C and 2F). Similarly, the image labeled as LD, but classified by the CNN model as ED, shows cells in the early stages of differentiation where cultures were sparser than usual and the cells lost their typical morphology (Fig. 2G). Among the images that were classified with equal confidence as ED and LD, the two top ones (Fig. 2B and 2E) are from the early stage of differentiation (day 11 and 14, respectively). One likely cause for a difficult classification of this stage is that day 14-cultures can display different morphologies depending on slight variations in cell density (slightly higher density in Fig. 2B, slightly lower in Fig. 2E). The image for day 16 (Fig. 2H) shows cells in the late stage of differentiation, but cultures at the late stage can appear slightly blurry without distinct cell borders, which makes the classification more difficult.

Stem cell differentiation is a continuous process, and it may be difficult to define an exact time point dividing cell differentiation stage into early and late differentiation. Hence, to further investigate the ability of the CNN model to properly classify the images, classification accuracy was assessed at each day separately (Table 1). Accuracy was 1.00 for images taken on day 1-8 and 17-23, except for day 22, which had 0.98 accuracy. On days 14 and 16, which we used as point of division between ED and LD, accuracy was reduced to 0.86. According to the differentiation protocol, the differentiation process switches from progenitor stage to hepatocyte maturation on day 14. Taking into consideration that changes in cell function and morphology are gradual, the lower accuracy around day 14 is not unexpected. In addition, differences in cell morphology on day 14 caused by slight variations in cell density may contribute to the lower accuracy (see explanation in previous paragraph). However, an accuracy of 0.86 implies that the CNN nonetheless can recognize morphological changes that occur within just two days.

**Table 1.** Classification accuracy by differentiation day on hPSC images in the independent test set.

| Differentiation stage | Days of differentiation | Accuracy |
| --- | --- | --- |
| Early | 1 | 1.00 |
|  | 2 | 1.00 |
|  | 3 | 1.00 |
|  | 4 | 1.00 |
|  | 6 | 1.00 |
|  | 7 | 1.00 |
|  | 8 | 1.00 |
|  | 9 | 0.92 |
|  | 10 | 1.00 |
|  | 11 | 0.97 |
|  | 14 | 0.86 |
| Late | 16 | 0.86 |
|  | 17 | 1.00 |
|  | 18 | 1.00 |
|  | 19 | 1.00 |
|  | 21 | 1.00 |
|  | 22 | 0.98 |
|  | 23 | 1.00 |

### Experimental validation of cell maturation during late differentiation

In practice, researchers rely on experimental assays to verify the functionality and differentiation stage of cell cultures. Four properties of cells are routinely measured to assess cell identity, namely cell morphology, gene expression, protein expression and cell functionality. To investigate the differentiation efficiency of hPSCs towards hepatocytes, typical established marker genes and proteins, and functional assays that define the maturity of the cells are available (Zhao et al., 2013). While the CNN was able to learn important morphological changes in microscopy images, we also wanted to experimentally characterize biological changes over the course of differentiation towards hPSC-HEP. Importantly, it was necessary to confirm that cell cultures in the late stage (corresponding to LD images) indeed have a mature hepatocyte phenotype. Therefore, experimental assays were carried out where expression of hepatic marker genes and proteins, albumin secretion and glycogen storage were assessed at several time points. Experimental results are shown in Fig. 3.

**Figure 3.**
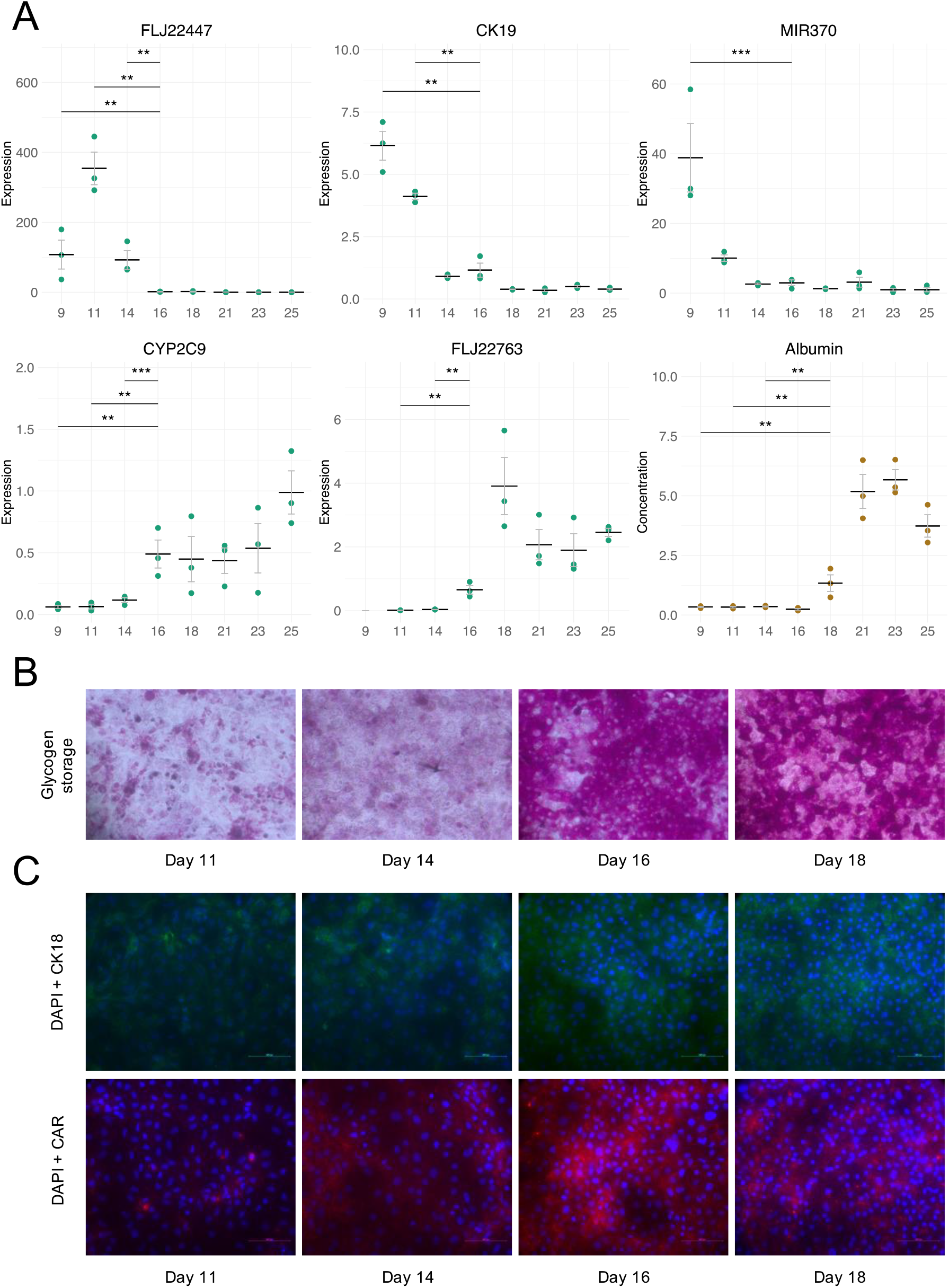
Experimental characterization of hPSCs undergoing differentiation towards hPSC-HEP. A: qPCR expression data for five markers of hepatocyte maturation (green) and albumin secretion (brown) in μg/mg protein/24h (N = 3, number of cell lines). Horizontal bar denotes the mean and error bars represent standard error. X axis shows number of days following differentiation initiation. Statistical testing was performed with repeated measures ANOVA followed the Tukey test. For clarity, only comparisons between day 9-14 and 16 (qPCR), and day 9-14 and 18 (albumin) are shown. No expression values were obtained for FLJ22763 day 9. ** denotes p < 0.01; *** denotes p < 0.001. B: Representative pictures of PAS-stained cultures visualizing glycogen storage on days 11-18. C: Representative pictures of immunocytochemical staining showing expression of CK18 and CAR on days 11-18.

The experimental results show clear changes in marker expression and cell function around days 14 to 16 after onset of differentiation. One of the key functions of the liver is metabolism of chemicals, where more than 90 % of reactions are catalyzed by the cytochrome P450 family of enzymes (Rendic and Guengerich, 2015). *CYP2C9*, one of the most abundant enzymes in adult liver, shows a significant upregulation on day 16 and expression is further elevated on day 25 (Fig. 3A). Other important liver functions include secretion of albumin and storage of glycogen. Albumin is a serum protein essential for maintenance of oncotic pressure, and is synthesized and secreted by hepatocytes (Buyl et al., 2015). The concentration of albumin in culture media was significantly elevated on day 18 and was further increased on day 21-25 (Fig. 3A). Another important liver feature is the storage of glycogen, which serves as a reservoir of glucose for other tissues in the body, and hepatic glycogen metabolism is important for maintenance of blood glucose levels (Bollen et al., 1998). The experimental data shows that the amount of glycogen stored in cell cultures was strongly increased on day 16 and onwards (Fig. 3B).

CK18 is a type-I intermediate filament protein highly concentrated in hepatocytes and cholangiocytes (epithelial cells of the bile duct), and comprises 5% of total liver protein (Uhlén et al., 2015). The constitutive androstane receptor (CAR), also known as NR1I3, is a member of the nuclear receptor superfamily (subfamily 1, group I, member 3) that is almost exclusively expressed in the liver. CAR is known to interact with key signaling pathways involved in drug, energy and bilirubin metabolism, and is an important biomarker for mature hepatocytes (Bae et al., 2021). The immunocytochemistry data shows an increasing expression of these two proteins in the late stages of the differentiation (day 16-18) (Fig. 3C).

These experimental results show that the hPSC cultures adopt hepatocyte features in the late stage of differentiation. We can also see that the CNN is able to classify images captured in the beginning and end of the differentiation period with accuracy close to 1.00 (Table 1). Thus, predictions made by the CNN based on cell culture morphology clearly reflect the underlying functional maturation of cells.

### Impact of image size, dataset size and data augmentation

In order to investigate the impact of image size on the training procedure, the CNN was trained with images of size of 400×400, 200×200 and 100×100 pixels. When comparing the model performance for images with resolution 400×400 and 200×200 pixels (Fig. 4A), the lower resolution resulted in a small reduction in accuracy and F1 during the validation of the model, while the computational time required for training was reduced 3.5-fold. Further reduction of image resolution to 100×100 pixels resulted in a drop of accuracy and F1 from 0.96-0.97 to around 0.91. Therefore, images of size 200×200 were used in this study.

**Figure 4.**
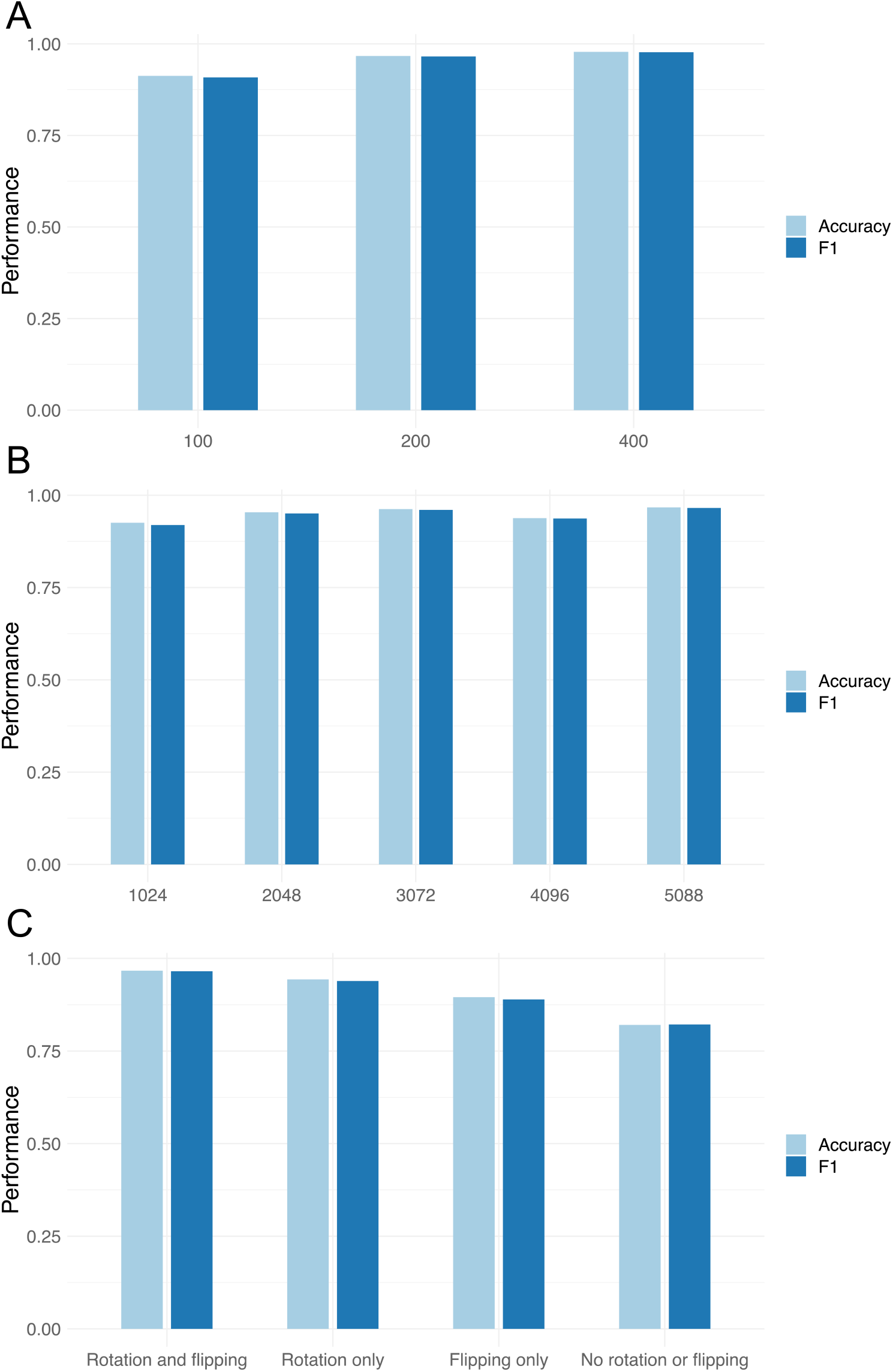
Impact of data processing on performance. Panels show impact of varying image size (A), dataset size (B) and image augmentation (C) on the validation accuracy and F1 of the CNN model.

One of the most common problems when training the CNN model is the lack of a sufficient number of images. It is difficult to estimate how much data is required for training since it is highly dependent on how challenging the classification problem is. Nonetheless, we investigated the sensitivity to changes sample size to provide an indication for the classification of early and late differentiated stem cells. Initially the set of 5,088 images was used for training, followed by a consecutive reduction in number of images until 1,024 images were left in the training dataset. Accuracy and F1 varied between 0.94 and 0.97 as the size of the dataset was reduced from 5,088 to 2,048 images (Fig. 4B). When only 1,024 images were used for training the model, a reduction in accuracy and F1 to 0.92-0.93 was observed.

Data augmentation was assessed in the present study by comparing classification performance on a dataset where images were flipped and rotated, with a dataset without one/both augmentation steps. The results in showed that both flipping and rotation indeed had a strong positive impact on performance of the CNN model by mitigating overfitting (Fig. 4C). Without augmentation, accuracy on the training set was close to 1.00 (data not shown), while accuracy on the validation set was only 0.82, which indicates that the model overfitted during the training. Excluding one step of image augmentation (flipping or rotation) likewise resulted in a reduction in classification accuracy.

## Discussion

Quality control of stem cell cultures is a crucial yet laborious process. Motivated by recent successes in AI-based microscopy image analysis (Lan et al., 2021; Liu et al., 2021; Zhu et al., 2021), we hypothesized that CNNs can be used to distinguish between different stages of hepatocyte differentiation, based on cell morphology in microscopy images. To evaluate this hypothesis, a CNN was trained to distinguish between images taken during early and late hepatocyte differentiation of hPSCs. Late differentiation was defined as the stage where hPSC-derived hepatic progenitor cells were differentiated further towards hPSC-derived hepatocytes (hPSC-HEP), which would correlate with morphological changes towards the functional, mature cell type. Our results showed that the CNN was highly successful at this task, achieving 0.96 accuracy on the independent test set, and close to accuracy of 1.00 on images taken in the beginning and end of the differentiation period, when differences in typical morphology were more pronounced. The predictions made by the CNN aligned well with experimental data on gene and protein expression, and functional features such as glycogen storage, reflecting the functional maturation of cells. In other words, the CNN output gives us important information about the cell maturation stage that in some cases may serve as an alternative to labor intensive and costly experimental assays. These results indicate that, with semi-automated CNN-based image analysis, it may be feasible to complement or even partially replace the need for extensive experimental characterization of differentiated hepatocytes. The adoption of computational methods to assess cell maturation could therefore simplify and accelerate stem cell production and research. Importantly, this would provide a less subjective method for morphological assessments than visual inspection performed by different trained researchers.

The results from this study showed a clear improvement of results when the image data was augmented. The assessment of data augmentation showed that the combination of rotating and flipping of images was the most successful approach, achieving a considerably higher accuracy compared to when these were not applied. Cutting the images into patches to increase dataset size also improved the performance of the model. Thus, the success of the CNN model trained in this study could therefore be largely attributed to the image augmentation and pre-processing steps. It is worth noting that we have used a fairly simple CNN model in this study, trained on approximately 1,300 images. This shows that even shallow CNNs can be successfully applied to classification of moderately sized microscopy datasets, depending on how challenging the classification problem is.

A strength of this study is the use of images from several cell lines obtained during everyday operation of a commercial lab. The original microscopy images were not specifically prepared or selected for the purpose of training a CNN. By using images obtained during everyday laboratory work, the data will be more closely representative of the heterogeneity due to e.g., instruments, lab personnel, reagent batches, variations in cell densities, etc. On the other hand, the images used to train, validate and test the CNN were generated by processing the original microscopy images. If the trained CNN was applied in practice, its task would be to classify cell cultures based on new uncropped microscopy images. The data used to train a CNN should be representative of those images, so the question arises to what extent training set image processing affects performance. One way to mitigate the risk of performance reduction in practice would be to use software that pre-processes the new microscopy images in real-time, generating multiple patches per image that the CNN classifies. This would result in several differentiation stage checks for different parts of the original image. Overall culture quality could then be assessed by combining the individual stage checks.

Other CNN architectures such as VGG16 (Simonyan and Zisserman, 2014) and ResNet (He et al., 2016) could also be investigated depending on how challenging the image classification task is. Given that our CNN achieved close to perfect accuracy, we did not explore this further. The accuracy was lower though for images captured on days near the change from progenitor medium to maturation medium. This is not unexpected, since morphological changes occur continuously during the differentiation, and no sharp distinctions can be observed at specific time points. Even when using molecular markers, it might be difficult to distinguish between cells around 14-16 days of culturing for the same reason. We do not believe this restricts the use of CNNs in future stem cell production and research though, as the cells would typically be assessed at the end of the differentiation process.

The results from this study clearly demonstrate the great potential of CNNs for addressing challenging image analysis problems related to stem cell culture characterization, where subtle differences may be of critical functional importance. However, this approach may also have great potential for quality control of stem cell products intended for regenerative medicine. In the promising area of advanced therapeutic medical products (ATMPs), the development of quality control procedures for assessment of cell identity and cell quality is a key challenge (Beheshtizadeh et al., 2022), for which CNNs may be a viable automated approach in large-scale cell production.

## Experimental procedures

### Image data

Convolutional neural networks (CNNs) were trained on phase-contrast microscope photographs of stem cell cultures captured at different time points during differentiation towards hPSC-HEP. Images of hPSCs differentiating into hepatocytes were obtained from Takara Bio Europe AB during routine differentiation processes, applying their current hepatocyte differentiation protocol (Cellartis iPS Cell to Hepatocyte Differentiation System, Cat. No. Y30055, Takara Bio Europe AB; see also section on Human Pluripotent Stem Cells Differentiation below), as well as a previous hepatocyte differentiation protocol also developed by Takara Bio Europe AB (Asplund et al., 2016).

A total of 1,331 images were collected from routine inspections of differentiation batches during a time period of several years. The images were obtained using a phase-contrast microscope (magnification 10x) (EclipseTi-U, Nikon, Amsterdam, The Netherlands), with an ANDOR Zyla sCMOS digital camera and then processed using NIS-Elements software package (version 4.30). Images were labeled as Early Differentiation (ED, days 1-14, in total 693 images) and Late Differentiation (LD, days 16-23, in total 638 images), respectively. The distinction between ED and LD was based on the change from differentiation to hepatocyte maturation culture medium on day 14.

### Pre-processing and augmentation

The dimensions of the original images were between 12801×024 and 2560×2160 pixels. To generate a larger number of images for the analysis, each image was cut into a maximum of six patches (1000×1000 pixels) using MATLAB. This was carried out for 70 % (932), 20 % (266) and 10 % (133) of the original images separately, and the three sets of patches were later used for training, validation and testing, respectively. The total number of patches generated was 6,972. Pixel values for all patches were scaled to values between 0 and 1.

Data augmentation was performed and the patches were downsized to 400×400, 200×200, and 100×100 pixels to investigate how image size affects the performance of the CNN. The patches were subsequently randomly flipped (vertically and horizontally) and rotated in the range between −0.2 and +0.2 of 360 degrees rotation with TensorFlow. This step was performed when the patches were given as input to the CNN. The flipped/rotated sets of patches are referred to as LargePatchAug, MediumPatchAug and SmallPatchAug, respectively, and each set contains 6,972 patches. These sets were used to evaluate how data augmentation affects CNN performance. The image processing and analysis workflow is illustrated in Fig. 5.

**Figure 5.**
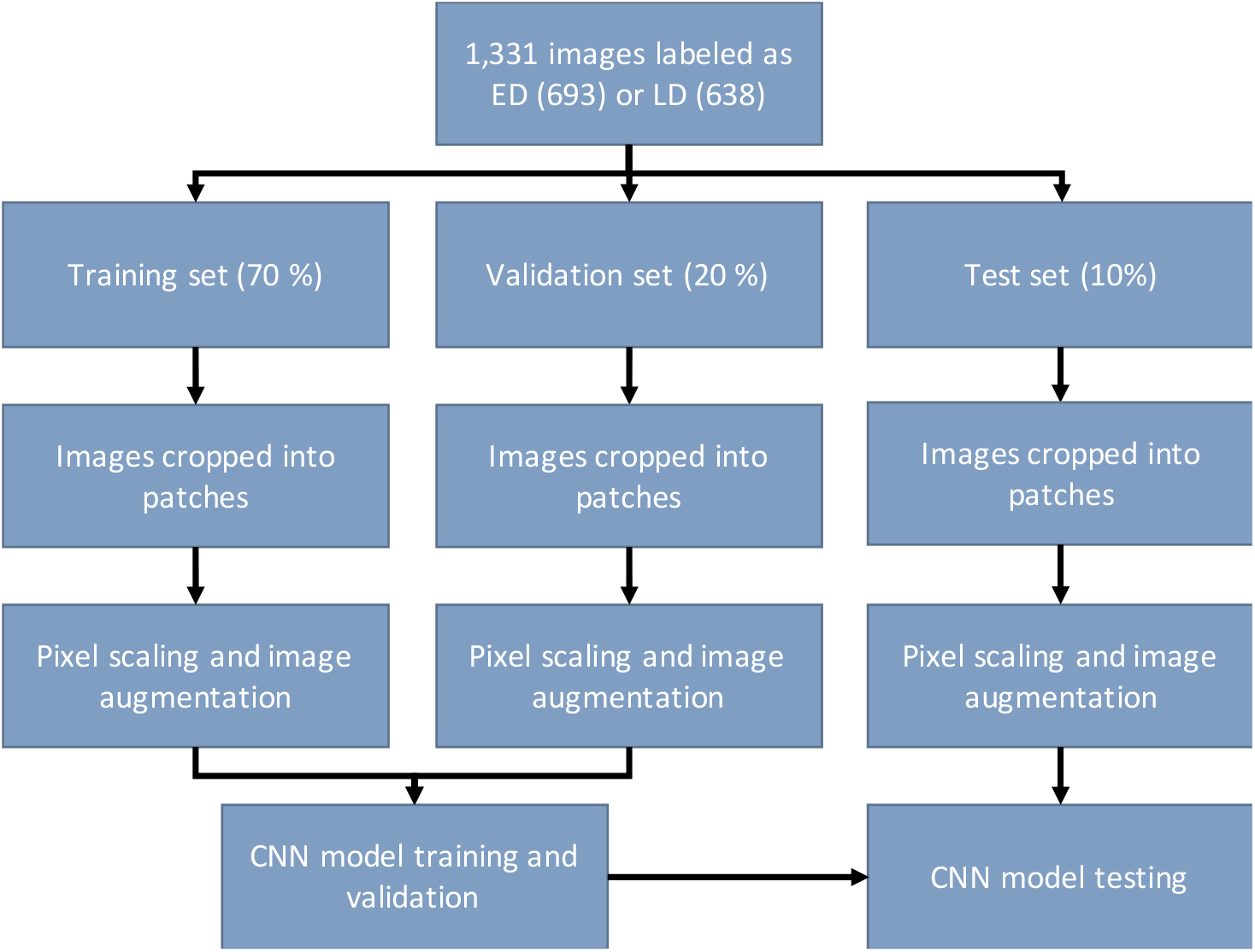
Image processing and analysis workflow. Abbreviations; ED: early differentiation, LD: late differentiation, CNN: convolutional neural network.

### Model specification

The CNN model was implemented in Python version 3.8.5 in the Anaconda 3 environment using Keras (Gulli and Pal, 2017) with TensorFlow 2.3.1 as a backend. The model architecture is illustrated in Fig. 6 and consists of an input layer followed by three sets of convolutional/ReLU/max-pooling layers. The last max-pooling layer is connected to a feed-forward network with one hidden layer and an output layer with two nodes for image classification (ED vs LD). The output layer used the Softmax activation function so the values of the output nodes sum to 1. This can be regarded as the probability of each image being ED or LD. Images were classified according to the class with highest probability.

**Figure 6.**
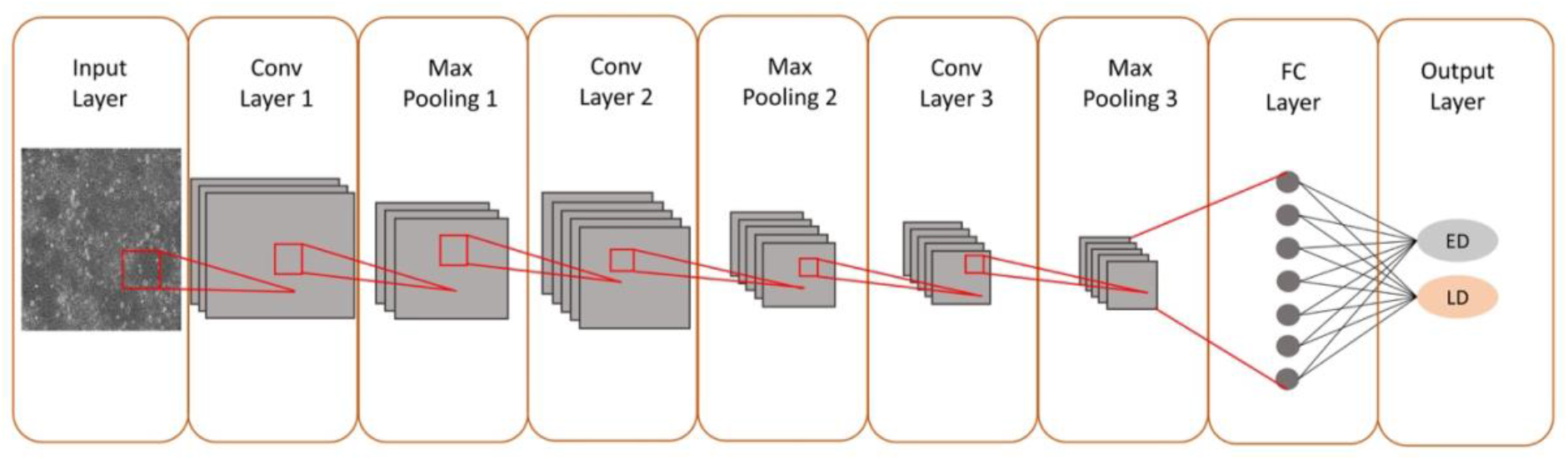
Architecture of the convolutional neural network used in this study. The network consists of three convolutional layers extracting features from an input image, three max pooling layers reducing the feature map size followed by two fully connected layers for image classification. The kernel, a filter extracting features from the image, is represented by a red color square.

### Training, validation and testing

The He Normal initializer was used to initialize the random weights of the CNN. The weights initialized by this method are random but also depend on the size of the upper layer of neurons in the network, allowing a faster and more efficient choice of weights range (He et al., 2016). According to Waisman et al. (Waisman et al., 2019), Adam, Adamax and Adagrad optimizers performed equally well when training CNNs for classification of mouse ESCs differentiating into epiblast-like cells. Therefore, Adam, an optimization algorithm designed especially for training deep neural networks (Kingma and Ba, 2014), was used in this study. The loss function used was binary cross entropy, since the network was optimized for binary classification of images. The model was fitted for 500 epochs and early stop with patience of 50 epochs. This allowed the model to stop training earlier if the loss function converges on a minimum value. Training was performed with a learning rate of 0.001 and mini batch size of 16.

The model was trained on 70 % of the LargePatchAug set and 20 % was used for validation/hyperparameter optimization. The remaining 10 % was reserved for testing. In addition, the following training set sizes were used to assess the impact of the number of training images on model performance: 4,096, 3,072, 2,048, 1,072. The number of images used for validation and testing was constant. Classification performance was reported using accuracy (ACC) and the F1 statistic according to Equation 1 and 2, respectively. True positives (TP) denote LD images classified as LD, false positives (FP) denote ED images classified as LD, true negatives (TN) denote ED images classified as ED, and false negatives (FN) denote LD images classified as ED.

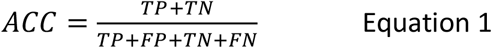

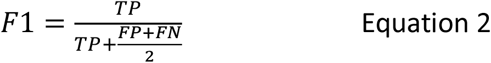

Analyses were performed on a personal computer with Windows 10 Pro 64 bits operating system, 3.3GHz Intel Xenon E-2126G CPU, 32 GB RAM, and NVIDIA Quadro P1000 GPU. GPU parameters were as follows: NVIDIA drivers 430.64, 512 Cuda cores, 4GB of dedicated video memory, and 20 GB of total available graphics memory. The runtime was between 20 min and 2h 15 min, depending on the dataset and hyperparameter settings.

### Human Pluripotent Stem Cells Differentiation

For the experimental validation of CNN predictions, hiPSC lines including Cellartis Human iPS Cell Line 22 (ChiPSC22) (Cat. No. Y00325), Cellartis Human iPS Cell Line 18 (ChiPSC18) (Cat. No. Y00305), and Cellartis Human iPS Cell Line 6b (ChiPSC6b), were obtained from Takara Bio Europe AB (Gothenburg, Sweden). The hiPSC lines were differentiated into human pluripotent stem cell derived hepatocytes (hiPSC-HEP) by using the Cellartis iPS Cell to Hepatocyte Differentiation System (Cat. No. Y30055, Takara Bio Europe AB) according to the manufacturer’s recommendations.

### RNA extraction, cDNA synthesis, and real time quantitative PCR

Cell samples were collected in RNAprotect Cell Reagent (Cat. No. 76526, QIAGEN) during the differentiation procedure at day 9, 11, 14, 16, 18, 21, 23, and 25. RNA was extracted from these samples using MagMAX-96 Total RNA Isolation Kit (Cat. No. AM1830, ThermoFisher). cDNA was synthesized applying High-Capacity cDNA Reverse Transcription Kit (Cat. No. 4368814, ThermoFisher). Real time quantitative PCR (RT-qPCR) was performed using TaqMan Fast Advanced Master Mix (Cat. No. 4444557, ThermoFisher) for the following markers: KRT19 (Hs00761767_s1, ThermoFisher), CYP2C9 (Hs04260376_m1, ThermoFisher), MIR370 (Hs04231551_s1, ThermoFisher), FLJ22447(Hs01382450_m1, ThermoFisher), and FLJ22763 (Hs01396927_m1). GAPDH (Hs99999905_m1, ThermoFisher) was used as a reference gene, and cDNA synthesized on a cocktail of RNA extracted from different cell types was used as calibrator (Asplund et al., 2016).

### Albumin Secretion Assay

Albumin secretion was determined on day 9, 11, 14, 16, 18, 21, 23, and 25 applying Albumin Human ELISA Kit (Cat.No. EHALB, ThermoFisher) on 24 h conditioned medium according to the manufacturer’s instruction. The results were normalized to total protein content. Protein content was measured applying Pierce BCA Protein Assay kit (Cat.No. 23227, ThermoFisher).

### Periodic Acid Schiff Staining

Periodic acid-Schiff Staining (PAS) was performed to detect glycogen storage in mature hiPSC-HEP. Fixed cells were incubated in periodic acid (Cat.No. 3951, SigmaAldrich) for 15 min and washed three times with dH_2_O. Then the cells were incubated in SCHIFF reagent (Cat.No. 3952016, SigmaAldrich) for 30 min. The cells were washed again three times with dH_2_O, and incubated in hematoxylin (Cat.No. GHS316, SigmaAldrich) for 90 sec. Finally, the cells were washed 3 times with dH_2_O.

### Immunocytochemistry

Cells were fixed at day 9, 11, 14, 16, 18, 21, 23, and 25 by incubation in 4% formaldehyde (Cat. No. 02176, Histolab) for 10 min at RT. The cells were permeabilized by incubation in 1% Triton X-100 (Cat.No. T8787, SigmaAldrich) in D-PBS +/+ (Cat.No. 14040-091, Gibco) for 15 min at RT, then the cells were blocked in 2% BSA (Cat.No. A9418, SigmaAldrich) in D-PBS +/+ for 60 min at RT. The cells were immunostained for the markers CK18 (Cat.No. MA5-12104, Invitrogen) and CAR (Cat.No. MA5-29208, Invitrogen). The antibodies were diluted in 0.1% BSA in D-PBS +/+ (1:100 for primary antibodies, and 1:1000 for secondary antibodies). The primary antibodies were incubated overnight at 4 °C. The secondary antibodies Goat anti-mouse IgG Alexa 488 (Cat.No. A11029, ThermoFisher), Donkey anti rabbit IgG Alexa 594 (Cat.No. A21207, ThermoFisher) and DAPI were incubated in the dark for 2 h at RT. The photographs were processed applying ImageJ software (http://imagej.nih.gov).

### Statistics

Data from qPCR and functional assays was imported into R version 4.1.2 (R Core Team, 2021) and log2-transformed prior to analysis. Statistical testing was performed with repeated measures ANOVA to identify statistically significant differences in marker gene expression and albumin/urea abundance between time points. Post-hoc testing was performed with Tukey’s test. Results were considered statistically significant where p < 0.05.

## Abbreviations

CNN: Convolutional neural network
AI: Artificial intelligence
iPSCs: Induced pluripotent stem cells
hiPSCs: Human induced pluripotent stem cells
mESCs: Mouse embryonic stem cells
hESCs: Human embryonic stem cells
PSCs: Pluripotent stem cells
hPSC: Human pluripotent stem cells
hPSC-HEP: hPSC-derived hepatocytes
ED: Early Differentiation
LD: Late Differentiation
qPCR: Quantitative polymerase chain reaction
ATMPs: Advanced therapeutic medical products

## Acknowledgements

This work was supported by Takara Bio Europe, Gothenburg, and by the University of Skövde, Skövde, under grants from the Knowledge foundation [20170302, 20200014].

## Author contributions

Conceptualization, B.U. and J.S.; Methodology, B.U. and K.M.S.; Software, K.M.S.; Validation, N.G.; Formal Analysis, B.U. and K.M.S.; Investigation, K.M.S., N.G. and B.K.M.; Resources, B.K.M.; Data Curation, K.M.S. and B.K.M.; Writing – Original Draft, K.M.S. and B.U.; Writing – Review & Editing, K.M.S., N.G., S.R.S., B.K.M., J.S. and B.U.; Visualization, K.M.S., S.R.S. and B.U.; Supervision, B.U.; Project Administration, B.U.; Funding Acquisition, B.U. and J.S.

